# Fractional Field Theory Approach to Protein Folding Dynamics

**DOI:** 10.1101/153999

**Authors:** Hosein Nasrolahpour

## Abstract

Understanding biological complexity is one of the most important scientific challenges nowadays. Protein folding is a complex process involving many interactions between the molecules. Fractional calculus is an effective modeling tool for complex systems and processes. In this work we have proposed a new fractional field theoretical approach to protein folding.

## 1. Introduction

Protein folding is a complex process involving many different interactions between the molecules that has attracted many attentions from physicist, chemists and biologists in recent years. Protein folding is the process by which proteins achieve rapidly and spontaneously their highly structured conformation with a certain biological function in a self-assemble manner, while misfolding process of protein can be seen as the failure to attain this fully functional conformation that may causes many different diseases such as: bone fragility, Alzheimer's disease, Parkinson disease and so on [1, 2]. There are many different approaches to address this issue such as: statistical mechanics and polymer dynamics etc. [3-7]. In the last decades, fractional calculus have found extensive applications in various fields of science from physics to biology, chemistry, engineering, economy and even in modeling of some human autoimmune diseases such as psoriasis[8-26]. Today fractional calculus is well known as an important effective modeling tool for complex systems and processes and can be used for describing various complex phenomena such as viscoelasticity, dielectric relaxations, fluid transport in fractal networks and so on [27-29].

The fractional variational principle can be considered as an important part of fractional calculus. Recently Agrawal has written a review article on this subject that can be found in [30] and discussed about various features of fractional variational calculus. Applications of fractional variational calculus have gained considerable popularity in science and engineering and many important results were obtained [31-39]. In our recent work we have propose the fractional sine-Gordon Lagrangian density, then using the fractional Euler-Lagrange equations, we have obtained fractional sine-Gordon equation [40]. Generalizing our previous results and using the approach present in [41, 42], we will propose a new fractional field theoretical approach to protein folding.

In the following, we will briefly review our mathematical tools. Then in Sec. 3 we present a new fractional protein Lagrangian density. Then using the fractional Euler-Lagrange equations we obtain its related equation of motion. Finally, in Sec. 4, we will present some conclusions.

## 2. Mathematical Tools

The fractional derivative has different definitions such as: Grünwald-Letnikov, Riemann-Liouville, Weyl, Riesz, Hadamard and Caputo fractional derivative [43], however in the papers cited above, the problems have been formulated mostly in terms of two types of fractional derivatives, namely Riemann-Liouville (RL) and Caputo. Among mathematicians, RL fractional derivatives have been popular largely because they are amenable to many mathematical manipulations. However, the RL derivative of a constant is not zero, and in many applications it requires fractional initial conditions which are generally not specified. Many believe that fractional initial conditions are not physical. In contrast, Caputo derivative of a constant is zero, and a fractional differential equation defined in terms of Caputo derivatives require standard boundary conditions. For these reasons, Caputo fractional derivatives have been popular among engineers and scientists. In this section we briefly present some fundamental definitions. The left and the right partial Riemann-Liouville and Caputo fractional derivatives of order *α_k_*, 0 <*α_k_* <1 of a function *f* depending on *n* variables, *x*_1_,…,*x_n_* defined over the domain 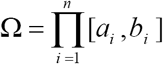 with respect to *x_k_* are as follow [35]:

The Left (Forward) RL fractional derivative

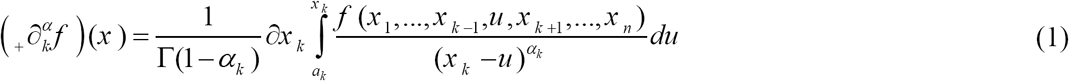

The Right (Backward) RL fractional derivative

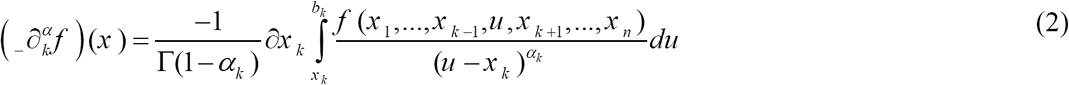

The Left (Forward) Caputo fractional derivative

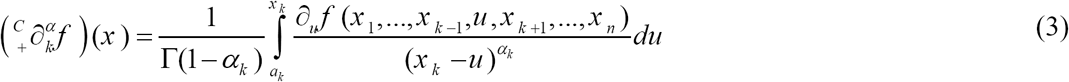

The Right (Backward) Caputo fractional derivative

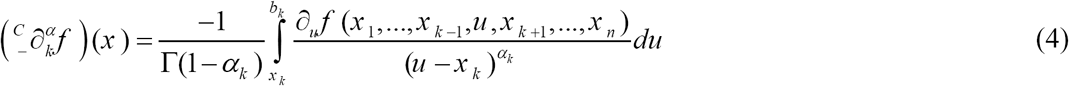

The fractional variational principle represents an important part of fractional calculus and has found many applications in physics. As it is mentioned in [30] there are several versions of fractional variational principles and fractional Euler-Lagrange equations due to the fact that we have several definitions for the fractional derivatives. In this work we use new approach presented in [35, 40] where authors developed the action principle for field systems described in terms of fractional derivatives, by use of a functional *S* (*ϕ*) as:

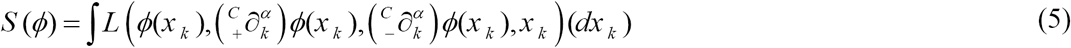

where 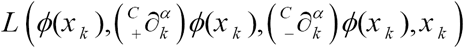 is a Lagrangian density function. Accordingly, *x_k_* represents *n* variables *x*_1_ to 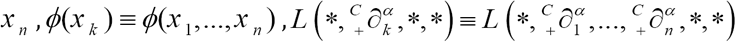, (*dx_k_*) ≡ *dx*_1_…*dx_n_* and the integration is taken over the entire domain Ω. From these definitions, we can obtain the fractional Euler-Lagrange equation as:

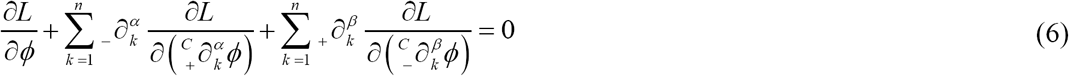

Above equation is the Euler-Lagrange equation for the fractional field system and fork *α, β* → 1, gives the usual Euler-Lagrange equations for classical fields.

## 3 Fractional Protein Lagrangian Density

Fractional dynamics is a field in theoretical and mathematical physics, studying the behavior of objects and systems that are described by using integrations and differentiation of fractional orders, i.e., by methods of fractional calculus [29]. Derivatives and integrals of non-integer orders are used to describe the behavior of nonlinear physical objects and systems that can be characterized by [29]:

i. power-law non-locality
ii. power-law long-term memory
iii. fractal-type property.

As an example in the realm of classical physics we can consider the well-known diffusion phenomena. The most known diffusion processes is the normal diffusion. This process is characterized by a linear increase of the mean squared distance:

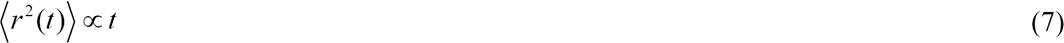

where *r* is the distance a particle has traveled in time *t* from its starting point. However there are many examples of phenomena in the natural sciences that violate this kind of behavior i.e. they are slower or faster than normal diffusion. In these cases (anomalous diffusions) the mean squared displacement is no longer linear in time:

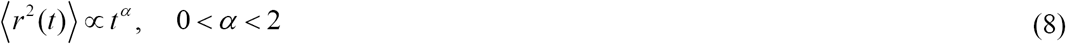

In recent years it is well known that generalization of the well-known diffusion equation and wave equation such that it includes derivatives of non-integer order with respect to time can describes phenomena that satisfy such a power law mean squared displacement.

Also it is well known that many biological systems are objects and systems with memory. As a result, the concept of fractional dynamics and in fact adopting fractional calculus can play an important role in the study of dynamical biological systems by taking advantage of the long memory properties of the fractional operators. In addition the advantage of modeling bio structures using fractional derivatives is the non-local property, and such these non-localities and memory effects in biological objects and systems mean that the next state of the system relies not only upon its present state but also upon all of its historical states [14].

Motivatetd by the above mentioned reasons in this section we present our new fractional Lagrangian model of protein folding that is in fact a fractional generalized version of the model presented recently in [41]. Using fractional Lagrangian model we will be able to consider complex nature of protein folding due to its memory effects and non-local nature. Following the model presented in [41, 42] we propose the protein Lagrangian including three terms:

I-Nonlinear unfolding *ϕ*^4^-protein at the initial state:

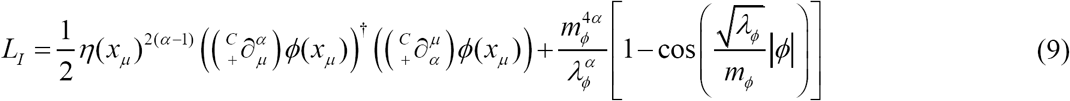

II-Nonlinear sources injected into the backbone, modeled by *Ψ*^4^ self-interaction:

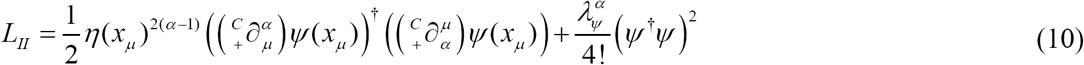

III-The interaction term (with the coupling constant Λ):

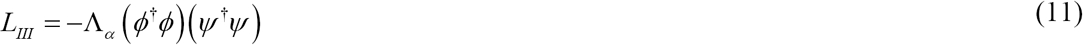

where *η*(*x_μ_*) is an arbitrary quantities with dimension of [second for *μ =* 0] and dimension of [meter for *μ =* 1,2,3] to ensure that all quantities have correct dimensions. The total potential (from all three terms) reads:

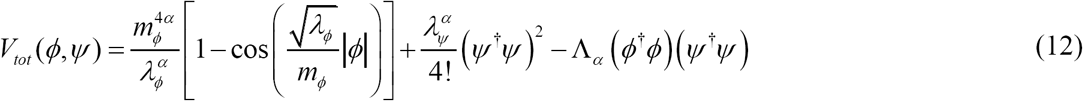

Assuming that *λ_ϕ_* is small enough to be approximately at the same order with *λ_ψ_*, the first term can be expanded in term of 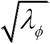, giving (up to the second order accuracy):

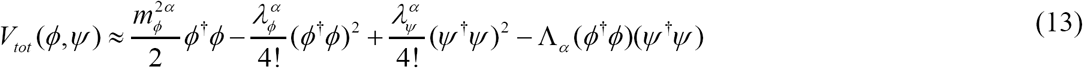

from which the total fractional Lagrangian: *L_tot_* = *L_I_ + L_II_ + L_III_* can be (up to the second order accuracy) approximated by:

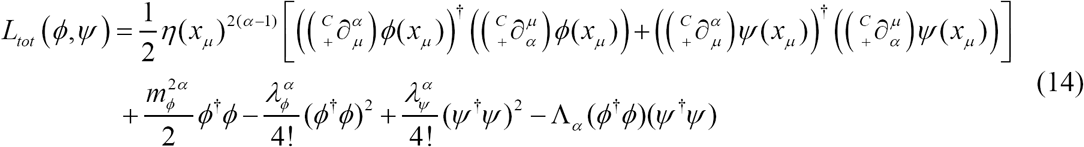

From the fractional Euler-Lagrangian equations for the total Lagrangian Eq. (6) we have: the following coupled and perturbed fractional Sine-Gordon equation and (nonlinear) fractional Klein-Gordon equation with cubic forcing in (1+1) dimension are derived:

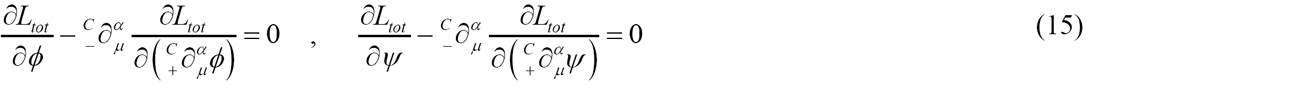

the following coupled and perturbed fractional Sine‑Gordon equation and (nonlinear) fractional Klein–Gordon equation with cubic forcing in (1+1) dimension are derived:

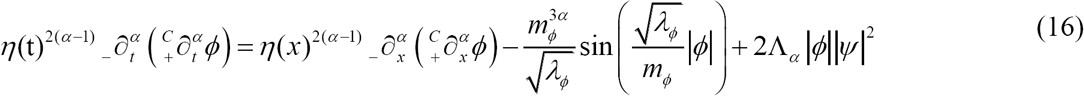

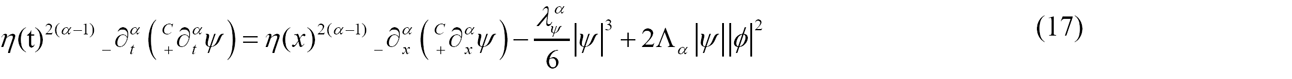

where *λ_ϕ_* and *λ_ψ_* ‑terms determine nonlinearities of backbone and source, respectively.

Solving these two coupled partial differential equations with specific boundary conditions would describe the contour of conformational changes for protein folding.

## 4. Conclusion

Fractional calculus is very useful tool for describing the behavior of nonlinear physical systems which are characterized by: power-law non-locality, power-law long-term memory and also fractal (or multifractal) properties. There exist many biological objects and systems with memory and nonlocal effects. In particular protein and its folding process has attracted many attentions from physicist, chemists and biologists in recent years. There are many different approaches addressing complex phenomena such as protein folding/misfolding however we believe that such these phenomena can be comprehensively understood by using fractional calculus and all of previous studies and models are only special cases of the model presented in this work. In this work we have proposed a new fractional field theoretical approach to protein folding. We have derived two coupled partial differential equations (i.e. fractional Sine–Gordon equation and fractional Klein‒Gordon equation) that their solutions with specific boundary conditions would describe the contour of conformational changes for protein folding. We hope to present our other result in future showing important role of fractional calculus in describing complex phenomena related to bio structures such as protein, DNA and RNA.

